# CBX4 regulates long-form thymic stromal lymphopoietin-mediated airway inflammation through SUMOylation in HDM-induced asthma

**DOI:** 10.1101/2021.05.24.445396

**Authors:** Shixiu Liang, Zicong Zhou, Zili Zhou, Jieyi Liu, Hangming Dong, Fei Zou, Haijin Zhao, Changhui Yu, Shaoxi Cai

**Affiliations:** Department of Respiratory and Critical Care Medicine, Chronic Airways Diseases Laboratory, Nanfang Hospital, Southern Medical University; Department of Occupational Health and Occupational Medicine, Guangdong Provincial Key Laboratory of Tropical Disease Research, School of Public Health, Southern Medical University, Guangzhou, Guangdong, China

**Keywords:** Asthma, airway inflammation, TSLP, SUMOylation, CBX4

## Abstract

**Rationale:** Thymic stromal lymphopoietin (TSLP) is present in two distinct isoforms, short-form (sfTSLP) and long-form (lfTSLP). lfTSLP promotes inflammation while sfTSLP inhibits inflammation in allergic asthma. However, little is known about the regulation of lfTSLP and sfTSLP during allergic attack in asthma airway epithelium.

**Methods and Results:** Here, we report that SUMOylation was enhanced in HDM-induced allergic asthma airway epithelium. Inhibition of SUMOylation significantly alleviated airway Th2 inflammation and lfTSLP expression. Mechanistically, CBX4, a SUMOylation E3 ligase, enhanced lfTSLP, but not sfTSLP, mRNA translation through the RNA binding protein, MEX-3B. MEX-3B promoted lfTSLP translation through binding of its KH domains to the lfTSLP mRNA. Furthermore, CBX4 regulated MEX-3B transcription in HBE through enhancing SUMOylation levels of the transcription factor, TFII-I.

**Conclusion:** We demonstrate an important mechanism whereby CBX4 promotes MEX-3B transcription through enhancing TFII-I SUMOylation, and MEX-3B enhances the expression of lfTSLP through binding to the lfTSLP mRNA and promoting its translation. Our findings uncover a novel target of CBX4 for therapeutic agents to lfTSLP-mediated asthma.

## Introduction

Thymic stromal lymphopoietin (TSLP) is an IL-7 like factor and has been reported to be an important epithelium-derived factor involved in the initiation and remodeling of allergic airway inflammation(***Adhikary, et al.,2020***). TSLP has been shown to contribute to T2-high inflammation by imparting its function on dendritic cells(***Ito, et al.,2005***), innate lymphoid cells(***Kim, et al.,2013***), and mast cells(***Astrakhan, et al.,2007***). Additionally, TSLP also been demonstrated to function in neutrophilic T2-low airway inflammation by activating dendritic cells to induce Th17 phenotype(***Tanaka, et al.,2009***). These data suggest that targeting TSLP might achieve broader effects. It has been reported that tezepelumab, an antibody targeting TSLP, significantly decreased the symptoms of patients with uncontrolled moderate-to-severe asthma exacerbations, irrespective of baseline blood eosinophils(***Corren, et al.,2017***). However, it has been demonstrated that TSLP exists in two distinct isoforms, long-form (lfTSLP) and short-form (sfTSLP), in human bronchial epithelial cells (only full length TSLP has been detected in mice)(***Harada, et al.,2009***). Previous data from our lab and others showed sfTSLP functions in antimicrobial activity and maintaining immune homeostasis, while lfTSLP promoted inflammation. In epithelium challenged with poly(I:C) and HDM, lfTSLP expression is upregulated, while sfTSLP expression is unaffected (***Dong, et al.,2016***; ***Harada, et al.,2009***). Given that sfTSLP are composed of 63 amino acid residues which are homologous to the lfTSLP C-terminal portion, the role of sfTSLP should be kept in mind when proposing therapeutic drug strategies to block lfTSLP in patients with asthma. Therefore, it is critical to understand the specific regulatory mechanisms of lfTSLP.

SUMOylation is an important post-translational modification implicated in many biological processes and diseases(***Yang, et al.,2017***). There are three mammalian SUMOylating enzymes (SUMO1–3). Activating (E1), conjugating (E2), and ligating (E3) enzymes are involved in protein SUMOylation(***Pichler, et al.,2017***). SUMOylation is removed by a family of SUMO specific proteases (SENPs)(***Kunz, et al.,2018***). Unlike polyubiquitylation, which facilitates protein degradation, SUMOylation regulates protein stability by affecting protein cellular localization and protein–protein or protein–DNA interactions(***Varejao, et al.,2020***). It has been reported that SUMOylation can regulate innate immunity and inflammatory responses through altering protein stability, such as SUMOylation of RIG-I and MDA5 by PIAS2 to increase their antivirus type I IFN responses(***Fu, et al.,2011***; ***Mi, et al.,2010***). In addition, SENP2 and SENP6 catalyze the de-SUMOylation of IRF3 and IKKγ, respectively, inhibiting TLR inflammatory responses and cellular antivirus responses(***Liu, et al.,2013***; ***Ran, et al.,2011***). These studies indicate SUMOylation might perform multiple functions in innate immunity and inflammatory responses via various substrates. It has been reported that the airway epithelium functions as an innate immunity barrier can activate by allergen through toll like receptor 3 and protease-activated receptor 2 and then cause an increase expression of TSLP(***Kato, et al.,2007***; ***Kouzaki, et al.,2009***). However, whether SUMOytion is involved in TSLP expression is unclear.

In the present study, we demonstrate that inhibition of SUMOylation alleviates airway inflammation and hyper-reactivity in an experimental model of allergic asthma. Furthermore, we identified CBX4, a SUMOylation E3 ligase, as playing a critical role in long form, but not short form, TSLP expression. Mechanistically, CBX4 regulates transcription of the RNA binding protein MEX-3B by enhancing transcription factor SUMOylation levels of TFII-I, resulting in enhanced expression of lfTSLP through MEX-3B binding to the lfTSLP mRNA and promoting its translation.

## Results

### Inhibition of SUMOylation Attenuates Airway Th2 Inflammation

To determine whether disproportional levels of SUMOylation/deSUMOylation occurs in asthma airway epithelium, we examined SUMO proteins level in an HDM-induced mouse model of asthma (Figure 1A). The immunohistochemical (IHC) results show a significant increase in expression of SUMO1 and SUMO2/3 in HDM-induced allergic airway epithelium (Figure 1B). Similar results were observed in mouse lung protein extracts (Figure 1C). These observations indicated enhanced levels of SUMOylation in asthma airway epithelium. Based on these observations, we wanted to address whether inhibition of SUMOylation may affect HDM-induced allergic airway inflammation. Mice were administered the SUMOylation inhibitor, 2-D08, before every challenge. In contrast to the only HDM-treated mice, HDM exposure induced much smaller peri-bronchial inflammation cells infiltration in the lungs of 2-D08-treated mice (Figure 1D). Mucus production also decreased in the airway of 2-D08-treated mice (Figure 1E). Consistent with these findings, leucocytes (eosinophils and neutrophils) (Figure 1F) and Th2 cytokines (IL-4, IL-5) (Figure 1G) from BALF, and total lgE (Figure 1H) in sera were reduced in 2-D08-treated mice. Airway hyper-reactivity (AHR) was measured after exposure to increasing doses of methacholine. We observed that 2-D08 treated mice significantly exhibited reduced AHR compared to HDM-treated mice (Figure 1I). Collectively, these results show that SUMOylation was enhanced after exposure to HDM, and that inhibition of SUMOylation can alleviate airway inflammation, mucus overproduction, and AHR.

**Figure 1.**
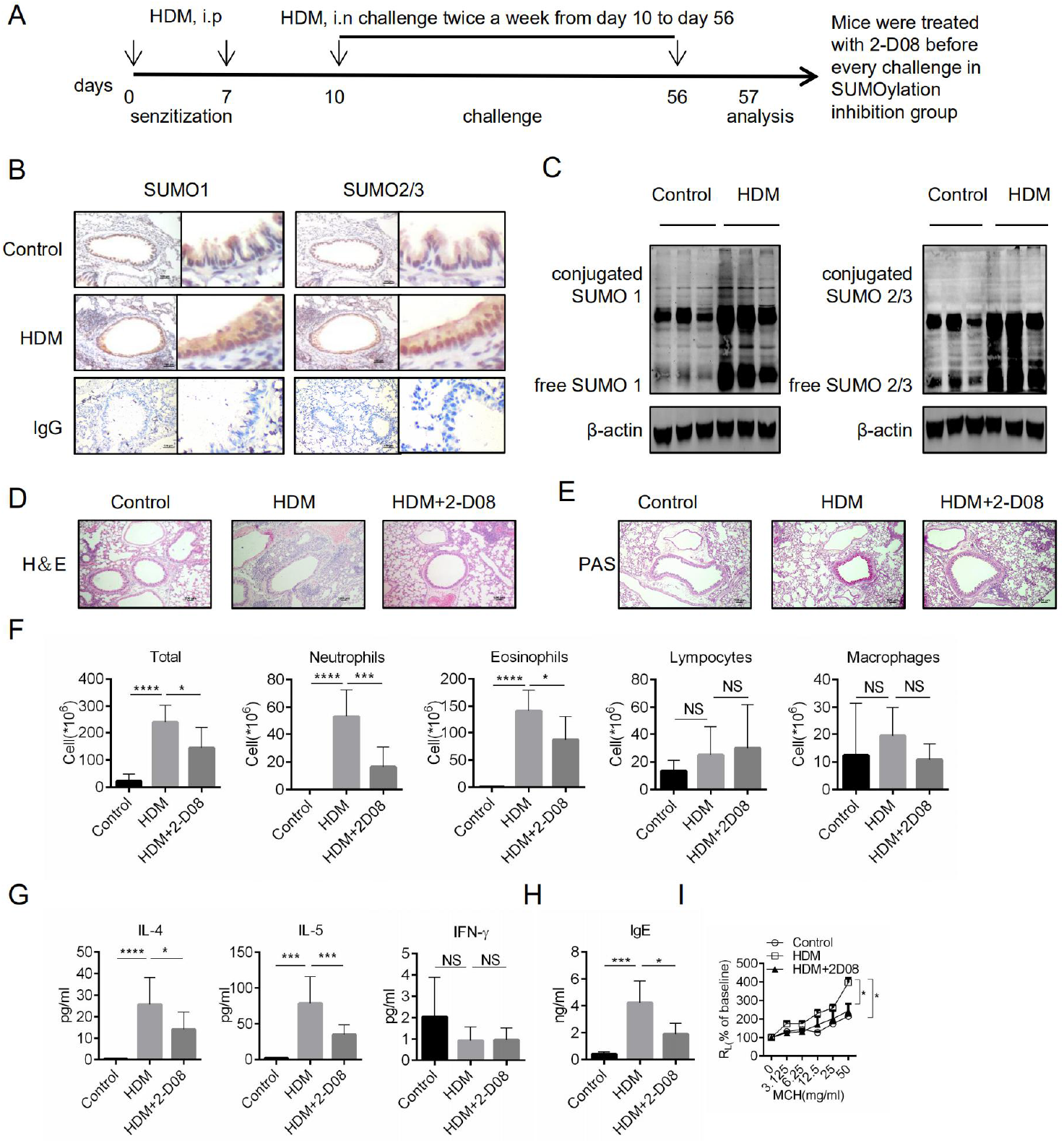
Inhibition of SUMOylation reduces HDM-induced allergic asthma. (A) Mice were sensitized (i.p) with 100 μl HDM or PBS on days 0 and 7, then challenged (i.n) with 100 μl HDM or PBS from day 10 to day 56, and samples were collected on day 57. Mice were treated with 2-D08 before every challenge in the SUMOylation inhibition group. (B) Lung sections were stained with SUMO 1 and SUMO 2/3 antibody by immunocytochemistry. (C) Immunoblot analysis of SUMO 1 and SUMO 2/3 in mice lung protein extract. Lung sections were stained with HE(D) and PAS(E) for assessing inflammation and mucus production. Quantification of inflammatory cell infiltration and airway mucus production in lungs was performed. (F) Cells in BALF were counted and classified following Wright-Giemsa staining. (G) Cytokines in BALF were measured by ELISA. (H) Serum lgE were qualified by ELISA. (I) Invasive measurement of dynamic airway resistance in response to increasing doses of methacholine. Data are representative of 2 independent experiments with at least 5 mice per group, and are presented as mean ± SEM. NS, Not significant. (D - G) One-way ANOVA with Bonferroni’s post hoc test was used. (H) Two-way ANOVA was used. \**p* < 0.05, \*\**p*< 0.01, \*\*\**p* < 0.001, and \*\*\*\**p*< 0.0001. Scale bar, 100 µm.

### lfTSLP Induction was Suppressed in 2-D08-Treated Asthma Mice and HBE

A study by Mitchell et al. reported that the airway epithelium is the main source of “alarmins” (e.g. IL-25, IL-33, and TSLP) in repose to allergens(***Mitchell and O’Byrne,2017***). Thus, we wondered whether inhibition of SUMOylation affects expression of these cytokines. We observed that IL-25 and TSLP, but not IL-33, proteins levels were reduced in 2-D08-treated mice lung extracts compared to HDM-treated mice (Figure 2A). To further investigate the effect of 2-D08 on the expression of “alarmins” in airway epithelium, we used lung histological sections stained for IL-25, IL-33 and TSLP. We found only TSLP expression was attenuated in airway epithelium of 2-D08 treated mice (Figure 2B-D). These results indicate that inhibition of SUMOylation decreases TSLP expression in the airway epithelium.

**Figure 2.**
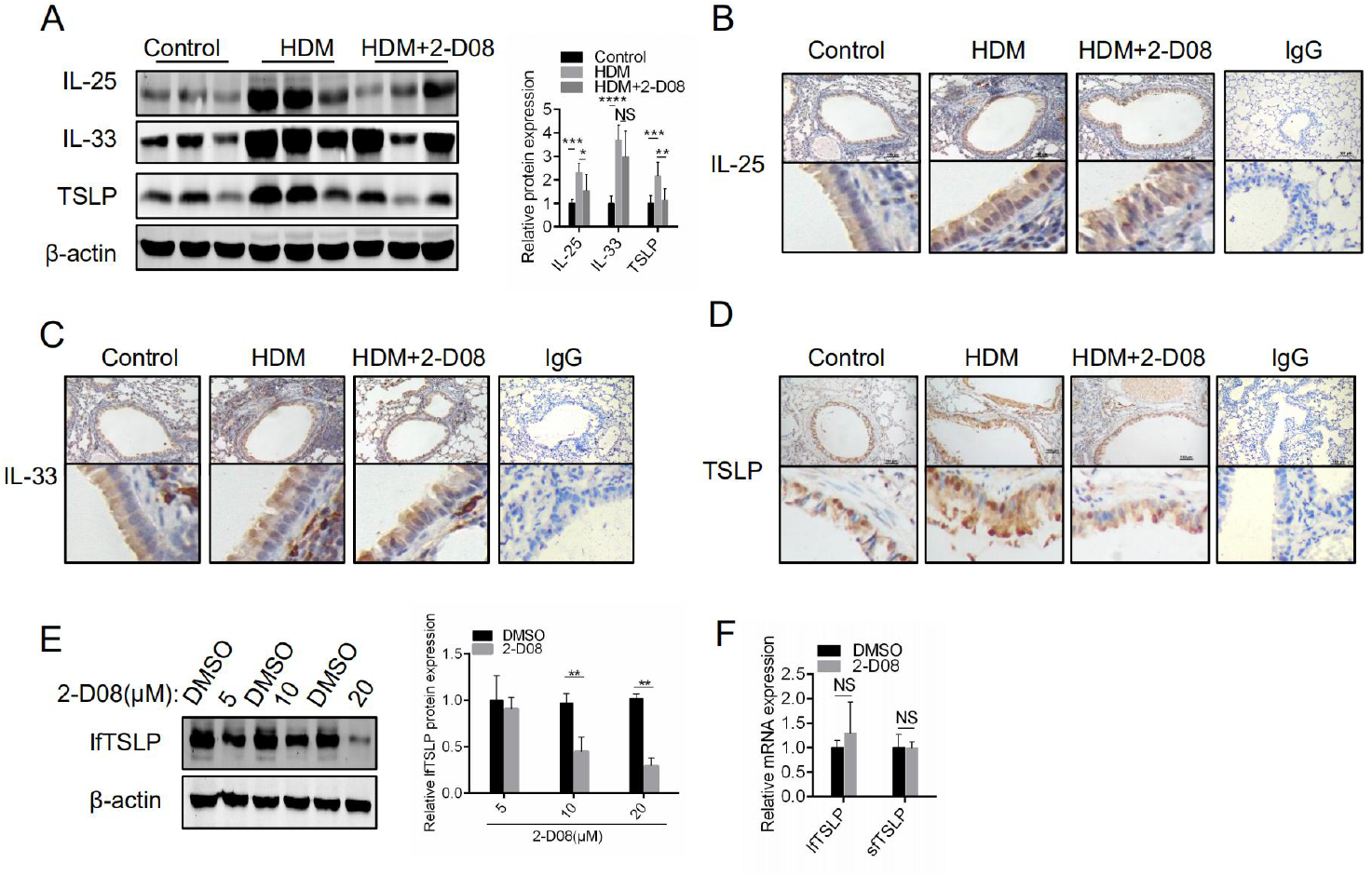
Inhibition of SUMOylation reduces lfTSLP protein expression. (A) Immunoblot analysis of IL-25, IL-33 and TSLP in mice lung protein extract. (B-D) Lung sections were stained with IL-25, IL-33 and TSLP antibody by immunocytochemistry. (E) lfTSLP protein levels were detected by Western blotting in HBE treated with different concentrations of 2-D08. (F) then analyzed lfTSLP and sfTSLP mRNA expression in HBE treated with 20 μM 2-D08 was measured by RT-PCR. Data are presented as mean ± SEM. Images show representative results for one of 3 or more experimental replicates. NS, Not significant. (A) One-way ANOVA with Bonferroni’s post hoc test was used. (E and F) Unpaired two tailed Student’s t-test was used. * *p* < 0.05, ** *p* < 0.01, and *** *p* < 0.001. Scale bar, 100 µm.

In humans, TSLP exists in two distinct isoforms, lfTSLP and sfTSLP, while only full length TSLP is found in mice(***Harada, et al.,2009***). We therefore investigated the effect of SUMOylation on lfTSLP and sfTSLP expression in human bronchial epithelial cells (HBE). We found that 2-D08-treated HBE display a significant reduction of lfTSLP protein expression (Figure 2E). Because there is no commercial primary antibody specific to sfTSLP, we designed primers specific to sfTSLP to measure the level of mRNA expression. We found that 2-D08 had no effect on sfTSLP expression (Figure 2F). Unexpectedly, 2-D08 also did not affect lfTSLP mRNA expression (Figure 2F). This finding suggests that SUMOylation may regulate lfTSLP post-transcriptionally. Altogether, these observations suggest that SUMOylation functions in lfTSLP expression in asthma model airway epithelium and HDM-treatment HBE.

### CBX4 is Involved in the Regulation of lfTSLP Expression in HBE

Given E3 enzymes and SENPs show specificity for substrates(***Pichler, et al.,2017***). For this reason, E3 enzymes and SENPs might be potential targets for therapeutic strategies(***Kumar and Zhang,2015***; ***Rabellino, et al.,2017***). Therefore, we performed RT-PCR to evaluate the expression of SUMOylation E3 ligases and deSUMOylation enzymes. Following exposure of HBE to HDM, we observed that the expression of SUMOylation E3 ligases are variably elevated while deSUMOylation enzymes SENPs are unaffected (Figure 3A). Among the upregulated SUMOylation E3 ligases, expression of CBX4 and PIAS1 increased significantly (Figure 3A and 3B). Consistent with the immunoblotting data in HBE in vitro, HDM-treated mice show a higher expression of CBX4 in lung extracts and airway epithelium compared to PBS-treated cells (Figure 3C and 3D). To evaluate the effect of CBX4 and PIAS1, HBE were transfected with siRNAs targeting CBX4 and PIAS1, respectively. We observed that siRNA knockdown of CBX4 resulted in a significant decrease in the level of lfTSLP protein while having no effect on IL-25 and IL-33 (Figure 3E and 3G). However, knockdown of PIAS1 did not affect the expression of lfTSLP, IL-25, or IL-33 (Figure 3F and 3H). Intriguingly, the level of lfTSLP mRNA expression was unaffected in HBE after knockdown of CBX4 or PIAS1 (Figure 3I and 3J). The discrepancy between the level of lfTSLP protein and mRNA indicated that CBX4 might regulate the expression of lfTSLP posttranscriptionally. Collectively, these data suggest that the SUMOylation E3 ligase CBX4 functions in lfTSLP protein expression in HDM-simulated HBE.

**Figure 3.**
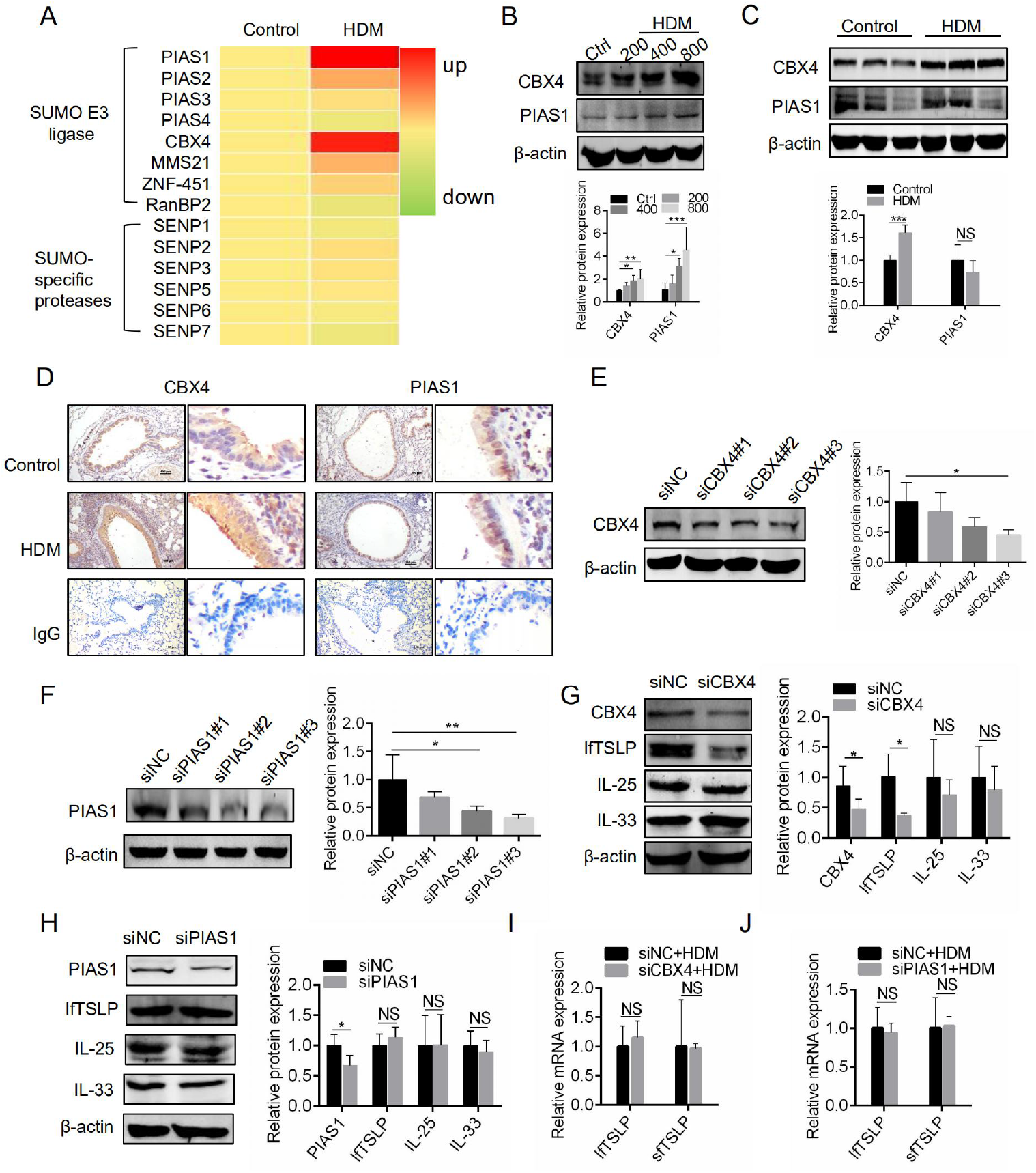
SUMOylation E3 ligase CBX4 regulates lfTSLP protein expression. (A) HBE were stimulated with 400 U HDM for 24 hr and expression of SUMOylation E3 ligases (PIAS1-4, CBX4, MMS21, ZNF451 and RanBP2) and deSUMOylation enzymes SENPs were measured by RT-PCR. (B and C) CBX4 and PIAS1 protein levels were measured by Western blotting in HDM-treated HBE (B) and mice lung protein extracts (C). (D) Lung sections were stained with CBX4 and PIAS1 antibody by immunocytochemistry. (E-H) Effects of CBX4 (G) and PIAS1(H) knockdown on IL-25, IL-33 and lfTSLP protein expression in HBE. (I and J) Effects of CBX4 (H) and PIAS1(I) knockdown on lfTSLP and sfTSLP mRNA expression. Data are presented as mean ± SEM. Images show representative results for one of 3 or more experimental replicates. NS, Not significant. (B, E and F) One-way ANOVA with Bonferroni’s post hoc test was used. (C, G, H, I and J) Unpaired two tailed Student’s t-test was used. * *p* < 0.05, ** *p* < 0.01, and *** *p* < 0.001. Scale bar, 100 µm.

### CBX4 is involved in the Regulation of lfTSLP Translation

In addition to its function as a SUMOylation E3 ligase, CBX4 is also a member of chromobox (CBX) protein family, which are canonical components of polycomb repressive complex 1 (PRC1) that functions as a transcription repressor(***Wotton and Merrill,2007***). The N-terminal chromodomain and two SUMO-interacting motifs (SIM 1/2) of CBX4 contribute to PRC1 and SUMO E3 ligase-dependent functions, respectively (Figure 4A). To investigate whether the effect of CBX4 in lfTSLP expression depends on its function in PRC1 or as a SUMO E3 ligase, expression of lfTSLP in HBE was measured following transfection with CBX4 plasmids bearing mutants in its chromodomain (CDM) or SIM (ΔSIM 1/2) (Figure 4A). we observed that ectopic expression of wild-type CBX4 (WT-CBX4) and CDM-CBX4 significantly increased the level of lfTSLP protein, whereas ΔSIM 1/2-CBX4 failed to do so (Figure 4B). This result suggests that CBX4 regulates lfTSLP through its SUMOylation function. This is finding agrees with the reduced level of lfTSLP protein observed in 2-D08-treated HBE (Figure 2C). In contrast, HBE treated with UNC3866,an inhibitor of the CBX4 chromodomain-histone interaction domain, did not affect lfTSLP expression compared to control (Figure 4C). Furthermore, these plasmid transfections did not affect lfTSLP mRNA expression in HBE (Figure 4D). These data indicate that SIM 1/2, but not the chromodomain of CBX4, are required for its regulation of lfTSLP expression.

**Figure 4.**
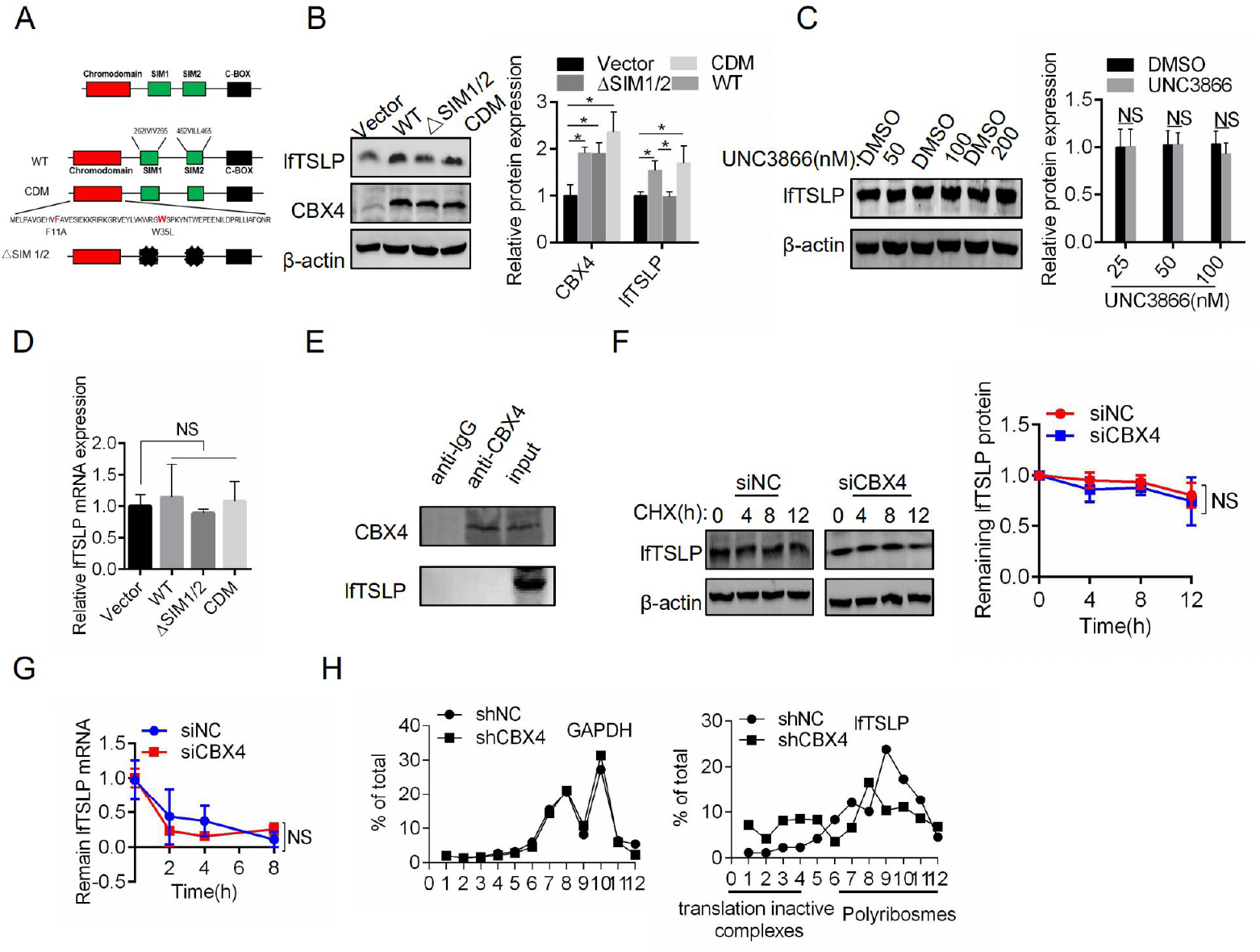
CBX4 regulates lfTSLP protein translation. (A) Schematic sketches of wild-type (WT) CBX4 and its mutants. (B) Immunoblots for the indicated proteins in HBE ectopically expressing the WT or CBX4 mutants for 48h. (C) HBE were treated with indicated concentrations of UNC3866 for 24hr. Expression of lfTSLP was determined by immunoblot analysis. (D) HBE were transfected with WT or CBX4 mutant plasmids for 48hr and lfTSLP mRNA expression was determined by RT-PCR. (E) HDM-treated HBE lysates were subjected to IP with anti-CBX4 antibody or anti-IgG antibody followed by immunoblot analysis with anti-lfTSLP antibody. (F) siNC or siCBX4 was transfected into HBE followed by treatment with 100 μg/mL cycloheximide (CHX) for the indicated amount of time. Expression of lfTSLP was determined by immunoblot analysis. (G) siNC or siCBX4 was transfected into HBE followed by treatment with 5 μg/mL actinomycin D for the indicated amount of time. The lfTSLP mRNA expression was measured by RT-PCR. (H) HBE transfected with shNC or shCBX4 were fractionated into cytoplasmic extracts through sucrose gradients. The distribution of lfTSLP and GAPDH mRNAs was quantified by RT-PCR analysis of RNA isolated from 12 gradient fractions. Data are presented as mean ± SEM. Images show representative results for one of 3 or more experimental replicates. NS, Not significant. (B) One-way ANOVA with Bonferroni’s post hoc test was used. (C) Unpaired two tailed Student’s t-test was used. (F and G) Two-way ANOVA was used. \**p* < 0.05.

As mentioned above, the effect of CBX4 on lfTSLP expression was SIM 1/2 dependent. To explore whether lfTSLP was regulated through CBX4-mediated SUMOylation directly, we performed immunoprecipitation to determine whether there is an interaction between CBX4 and lfTSLP. Unexpectedly, no interaction between these two proteins was observed (Figure 4E). Furthermore, HBE was treated with the protein synthesis inhibitor, cycloheximide (CHX), with or without CBX4 knockdown. The rate of degradation of lfTSLP protein was similar in HBE transfected with CBX4 or negativesiRNA (Figure 4F). These results indicate that CBX4 did not regulate lfTSLP expression post-translationally. We therefore speculated that CBX4 might affect mRNA translation of lfTSLP. As mentioned above, CBX4 knockdown in HBE results in reduced levels of lfTSLP protein while the level of mRNA was un affected. Therefore, we treated HBE with the RNA polymerase II inhibitor actinomycin D for different time intervals with or without CBX4 knockdown. We observed that knockdown of CBX4 did not affect lfTSLP mRNA degradation in HBE compared to the control group (Figure 4G). Furthermore, to determine whether CBX4 affects lfTSLP translation, we performed polysome fractionation to analyze lfTSLP mRNA distribution profiles through sucrose gradients to separate ribosomal subunits (40S and 60S), monosomes (80S), and progressively larger polysomes in HBE subjected to CBX4 siRNA transfection. Each group was divided into 12 fractions and the levels of lfTSLP and GAPDH mRNA were assessed by RT-PCR analysis. We observed that lfTSLP mRNA shifted from fractions enriched for translating polyribosomes fractions (5–12), indicative of enhanced translation, to fractions containing translation-dormant complexes, including mRNPs, ribosome subunits, and monosomes (fractions 1–4) after CBX4 knockdown in HBE (Figure 4H). Overall, these results support the idea that CBX4 enhances lfTSLP translation without affecting its mRNA stability in HDM-treated HBE.

### CBX4 Promotes lfTSLP Translation through the RNA Binding Protein MEX-3B

It has been reported that RNA binding proteins (RBPs) are essential for posttranscriptional gene regulation, linking RNA transcription, splicing, export, rate of translation, and stability(***Licht and Jantsch,2016***; ***Wang, et al.,2018***). In all of these processes, RBPs coordinate the regulation of the amount of proteins produced from mRNA transcripts. To determine whether CBX4 regulates lfTSLP translation through RBPs, HBE transfected with CBX4 siRNA and control siRNA were subjected to whole-transcriptome sequencing. Sequencing analysis revealed that a total of 283 genes (197 upregulated and 86 downregulated) significantly changed in HBE transfected with siRNA targeting CBX4 compared to control. Included in the 86 downregulated genes were MEX-3B and RBM44 (Figure 5A), two RBPs that play an important role in posttranscriptional gene regulation and are involved in a variety of diseases(***Oda, et al.,2018***; ***Yamazumi, et al.,2016***). Interestingly, RT-PCR analysis confirmed that MEX-3B, but not RBM44, is regulated by CBX4 (Figure 5B). Additionally, MEX-3B protein expression decreased after CBX4 knockdown in HBE (Figure 5C). Furthermore, we observed that MEX-3B was upregulated in HDM-stimulated HBE (Figure 5D-E).

**Figure 5.**
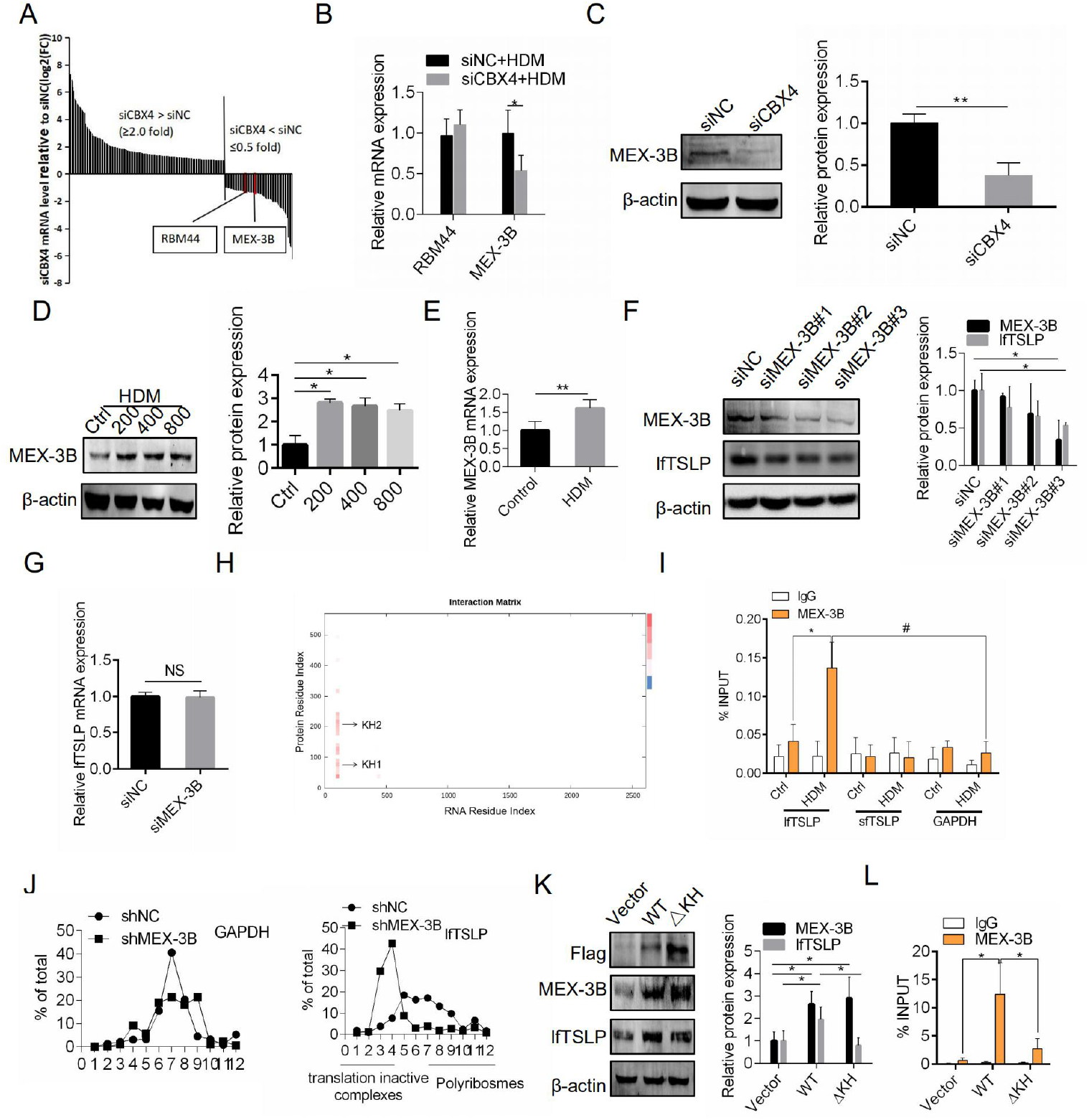
CBX4 regulates lfTSLP protein translation through MEX-3B. (A) HBE were transfected with siNC or siCBX4 and subjected to RNA-seq. There were 197 upregulated and 86 downregulated genes. Among the 86 downregulated genes, there were two RNA binding proteins: MEX-3B and RBM44. (B) Expression of MEX-3B and RBM44 was validated by RT-PCR in HBE transfected with siNC or siCBX4. (C) HBE were transfected with siNC or siCBX4. MEX-3B protein expression was determined by immunoblot analysis. (D and E) HBE were stimulated with HDM and MEX-3B protein levels and mRNA were measured by immunoblot analysis and RT-PCR, respectively. (F and G) siNC or siMEX-3B was transfected into HBE. Expression of lfTSLP protein and mRNA were measured by immunoblot analysis and RT-PCR, respectively. (H) The potential binding sites of MEX-3B protein and lfTSLP mRNA were predicted by the catRAPID database. (I) RT-PCR analysis of lfTSLP and sfTSLP mRNA that co-immunoprecipitated with mouse immunoglobulin G (IgG) or anti-Mex-3B antibody in HBE. GAPDH mRNA was used as a negative control. (J) HBE transfected with shNC or shMEX-3B were fractionated into cellextracts through sucrose gradients. The distribution of lfTSLP and GAPDH mRNAs was quantified by RT-PCR analysis of RNA isolated from 12 gradient fractions. (K) Immunoblots for the indicated proteins in HBE transfected with control vector, Flag-tagged Mex-3B, or Flag-tagged Mex-3B-mutKH. (L) RT-PCR analysis of lfTSLP mRNA co-immunoprecipitated with mouse immunoglobulin G (IgG) or anti-Mex-3B antibody in HBE transfected with control vector, Flag-tagged Mex-3B, or Flag-tagged Mex-3B-mutKH. Data are presented as mean ± SEM. Images show representative results for one of 3 or more experimental replicates. NS, Not significant. (B, C, E and G) Unpaired two tailed Student’s t-test was used. (D, F, I, K and L) One-way ANOVA with Bonferroni’s post hoc test was used. \**p* < 0.05 and ** *p* < 0.01.

MEX-3B is a member of MEX-3 family, which comprises MEX-3A, MEX-3B, MEX-3C, and MEX-3D. It has been reported that the MEX-3 family of proteins bind to specific mRNAs and regulate the expression of their proteins depending on two K-homology (KH)-type RNA-recognition domains(***Buchet-Poyau, et al.,2007***). Therefore, we also explored whether other MEX-3 family proteins were regulated by CBX4, and found that CBX4 knockdown had no effect on MEX-3A, MEX-3C, and MEX-3D expression (Figure S1). However, whether MEX-3B is involved in the posttranscription regulation of lfTSLP remains unclear. To determine if MEX-3B is involved in the posttranscription regulation of lfTSLP, lfTSLP protein and mRNA levels were measured in HBE transfected with siRNA targeting MEX-3B. We observed that lfTSLP protein expression, but not mRNA expression, significantly decreased after MEX-3B knockdown (Figure 5F and 5G). To address whether MEX-3B exerts its effect through binding to lfTSLP mRNA, we predicted the potential binding sites between the MEX-3B protein and lfTSLP mRNA using the protein-RNA interaction database, catRAPID. The results suggest that MEX-3B may interact with the lfTSLP mRNA 5’UTR through its KH domains (Figure 5H). To confirm this, lysates from HDM-treated HBE were subjected to immunoprecipitation with a MEX-3B primary antibody, then lfTSLP mRNA associated with complexity was detected by RT-PCR. We observed that the MEX-3B protein interacted with the lfTSLP mRNA, but not the sfTSLP mRNA, and that the interaction was enhanced upon HDM stimulation (Figure 5I). Furthermore, knockdown of MEX-3B repressed lfTSLP translation (Figure 5J). To explore whether MEX-3B facilitating lfTSLP mRNA translation depends on its KH domains, HBE were transfected with plasmids encoding wild-type MEX-3B or a KH domains mutant MEX-3B. We found that wild-type MEX-3B transfection significantly increased lfTSLP protein levels while the KH domains mutant failed to do so (Figure 5K). Additionally, the MEX-3B KH domains mutant exhibited a lower level of association with the lfTSLP mRNA compared to wild-type MEX-3B (Figure 5L). These data suggest that CBX4 promotes lfTSLP translation through the RNA binding protein MEX-3B, and that MEX-3B binds to the lfTSLP mRNA and facilitates its translation through its KH domains.

### The Transcription Factor TFII-I Binds the MEX-3B Promoter

As mentioned above, CBX4 regulates MEX-3B mRNA and protein expression. Additionally, CBX4 knockdown in HBE treated with actinomycin D did not affect lfTSLP mRNA degradation compared to the control group (Figure 6A). This suggested that CBX4 may regulate the transcription of MEX-3B. Therefore, we speculated that CBX4 may directly bind to the MEX-3B promoter and scanned the potential binding sites between CBX4 and the MEX-3B promoter through hTFtarget database. Unfortunately, no potential binding sites were found on the MEX-3B promoter (Figure S2). Furthermore, we found that the CBX4-histone interaction inhibitor UNC3866 failed to alter MEX-3B expression (Figure 6B). These data suggested that CBX4 might not function as a transcription factor for MEX-3B by binding directly to the promoter.

**Figure 6.**
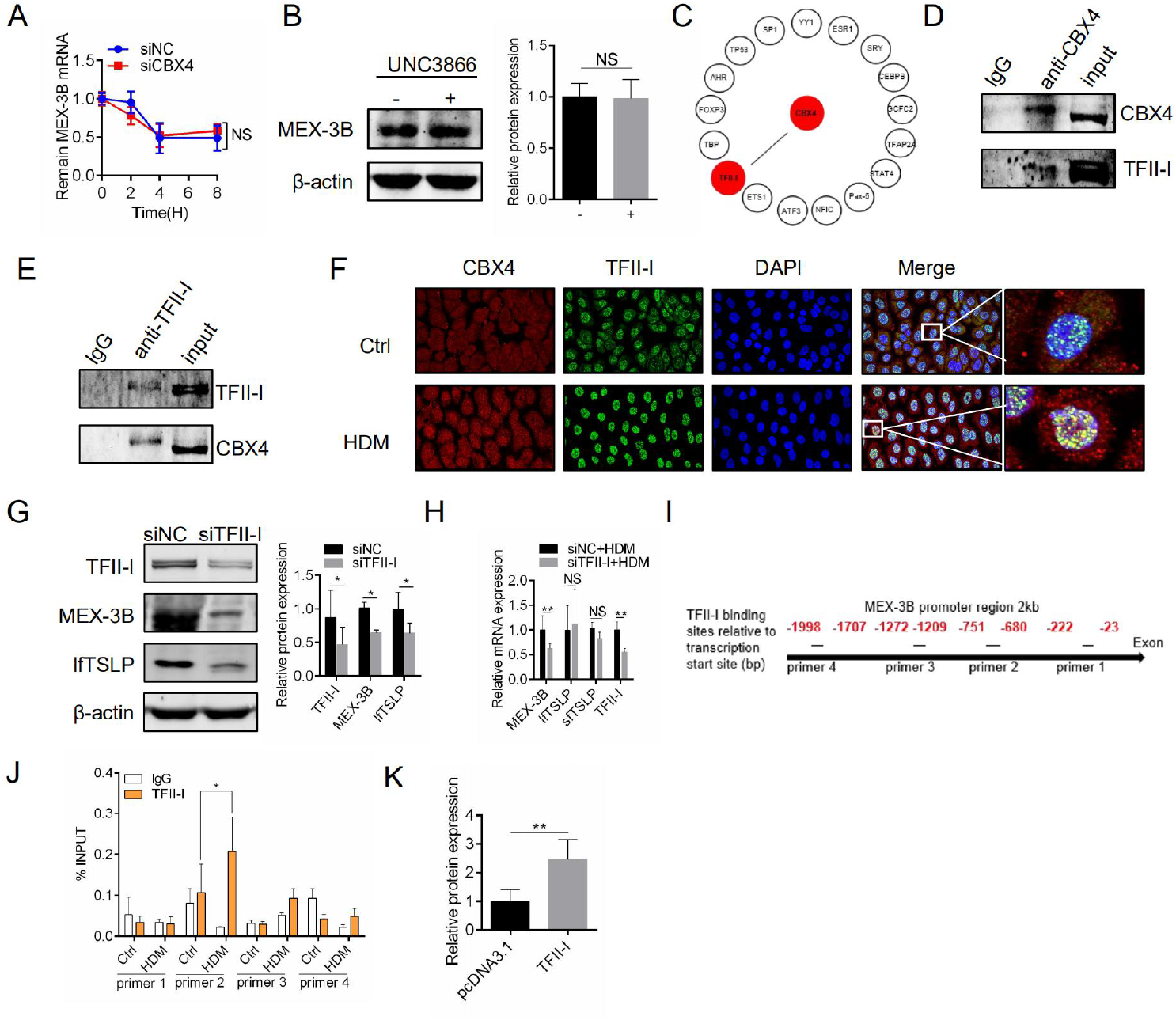
TFII-I is a transcriptional activator of MEX-3B. (A) siNC or siCBX4 was transfected into HBE followed by treatment with 5 μg/mL actinomycin D for the indicated amount of time. MEX-3B mRNA expression was measured by RT-PCR. (B) HBE were treated with the indicated concentrations of UNC3866 200 nM for 24hr. The MEX-3B protein level was determined by immunoblot analysis. (C) Potential transcription factors were scanned by the ALGGEN database. Prediction of an interaction between CBX4 and these factors were conducted by GeneMANIA. (D) HBE lysates were subjected to IP with anti-CBX4 antibody or anti-IgG antibody followed by immunoblot analysis with anti-CBX4 and TFII-I antibody. (E) HBE lysates were subjected to IP with anti-TFII-I antibody or anti-IgG antibody followed by immunoblot analysis with anti-TFII-I and CBX4 antibody. (F) Co-localization of CBX4 and TFII-I was analyzed by immunostaining of HBE with anti-CBX4 and TFII-I via confocal microscopy. (G) Immunoblots for the indicated proteins in HBE transfected with siNC or siCBX4. (H) RT-PCR for the indicated mRNA expression levels in HBE transfected with siNC or siCBX4. (I) Localization of TFII-I-binding sites in MEX-3B promoter. (J) HBE were treated with HDM 400 U 24 hr followed by ChIP with anti-TFII-I antibody or nonrelated IgG. Precipitated DNAs were quantified by RT-PCR using five MEX-3B promoter-specific primers covering five TFII-I-binding sites. (K) Human 293T cells transfected with pcDNA3.1(+)-TFII-I together with firefly luciferase reporter and pRL-tk-renilla plasmids for 24 hr. Data are presented as mean ± SEM. Images show representative results for one of 3 or more experimental replicates. NS, Not significant. (A) Two-way ANOVA was used. (B, G, H and K) Unpaired two tailed Student’s t-test was used. (J) One-way ANOVA with Bonferroni’s post hoc test was used. \**p* < 0.05 and ** *p* < 0.01.

Therefore, we hypothesized that CBX4 might function as a transcriptional coactivator of MEX-3B. To confirm this, we first used the ALGGEN database to predict the transcription factors that might bind to the promoter of MEX-3B. The result show that there are 17 potential transcription factors of MEX-3B (Figure S3). Next, we conducted an interaction prediction between CBX4 and these factors through GENEMANIA. Intriguingly, only general transcription factor II (TFII-I, encoded by GTF2I) may interact with CBX4 (Figure 6C) and immunoprecipitations confirmed the interaction (Figure 6D and 6E). Similarly, colocalization of CBX4 and TFII-I was also observed upon HDM stimulation in HBE (Figure 6F). Next, we investigated whether TFII-I could regulate MEX-3B expression. Knockdown of TFII-I resulted in a significant decrease of MEX-3B protein and mRNA levels (Figure 6G and 6H). To validate TFII-I as a transcription factor of MEX-3B, HBE treated with HDM were subjected to chromatin immunoprecipitation (ChIP). Because the database indicated five major potential TFII-I binding sites in the MEX-3B promoter region, five MEX-3B promoter-specific primers covering these sites were designed for RT-PCR. The results of the ChIP assay identified binding of TFII-I to the MEX-3B promoter (−680∼-751), which was significantly enhanced in HBE simulated by HDM (Figure 6J). Furthermore, we observed that overexpression of TFII-I promoted transcriptional activity of MEX-3B using a luciferase reporter assay (Figure 6K). These results suggest that TFII-I is a transcriptional activator of MEX-3B.

### CBX4-mediated TFII-I SUMOylation Enhanced the Transcription of MEX-3B

Previous studies have demonstrated that the transcriptional activity of TFII-I could be enhanced by SUMOylation(***Tussie-Luna, et al.,2002***). Therefore, we suspected that CBX4 might alter the level of TFII-I SUMOylation and, therefore, transcriptional activity. We observed that CBX4 knockdown caused a significant reduction of TFII-I binding to MEX-3B promoter and transcriptional activity (Figure 7A and 7B). Interestingly, HDM stimulation or CBX4 knockdown did not alter TFII-I expression in HBE (Figure 7C). Consistently, the expression levels of TFII-I were equal in control, HDM-treated, and 2-D08-treated mice airway epithelium (Figure 7D). These results indicated that CBX4 regulated only TFII-I transcriptional activity, not its expression. To confirm that CBX4 regulates TFII-I transcriptional activity through SUMOylation, the level of TFII-I SUMOylation was measured by co-immunoprecipitation in HBE stimulated with HDM. We found the conjunction of TFII-I and SUMO1 was enhanced after exposuring to HDM while decreasing after knock down CBX4 in HBE (Figure 7E and 7F). Similarly, colocalization of TFII-I and SUMO1 increased upon HDM simulation, and decreased upon CBX4 knockdown (Figure 7G and H). Next, we investigated whether CBX4 regulation of the transcription activity of TFII-I depends on the SIM structure of CBX4. For convenience of transfection, human 293T cells were transfected with plasmids expressing TFII-I and different mutant forms CBX4. CHIP and luciferase reporter gene assay showed that overexpression of CBX4 promoted TFII-I binding to the MEX-3B promoter and transcription activity, and this effect persisted with the transfection of CDM-CBX4, but not ΔSIM 1/2-CBX4 (Figure 7G and 7H). These observations support the hypothesis that CBX4 increases TFII-I SUMOylation and enhances the binding of TFII-I to the MEX-3B promoter, resulting in an increase in TFII-I-mediated MEX-3B transcription.

**Figure 7.**
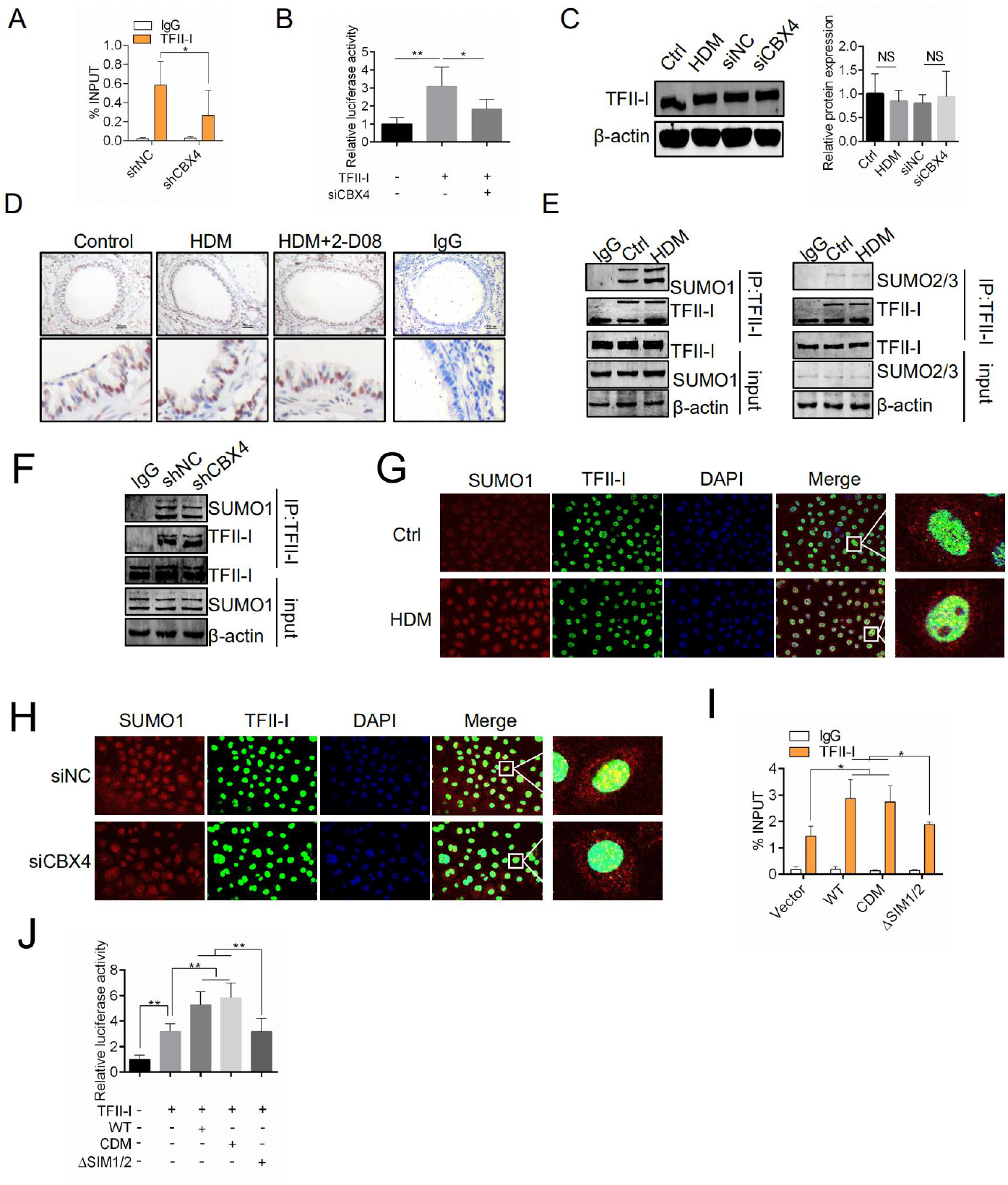
CBX4 regulates TFII-I transcriptional activity through SUMOylation. (A) HBE were transfected with shNC or shCBX4 followed by ChIP with anti-TFII-I antibody or IgG. Precipitated DNAs were quantified by RT-PCR using MEX-3B promoter-specific primers covering TFII-I-binding sites. (B) Human 293T cells transfected with pcDNA3.1(+)-TFII-I and siCBX4, together with firefly luciferase reporter and pRL-tk-renilla plasmids for 24 hr. (C) Immunoblot analysis of TFII-I protein levels in HBE treated with HDM, siNC, or siCBX4. (D) Lung sections were stained with TFII-I antibody by immunocytochemistry. (E) Enhanced TFII-I SUMOylation in HDM-treated HBE. HDM-treated HBE was subjected to IP with anti-TFII-I followed by immunoblot analysis with anti-SUMO 1 and SUMO 2/3 antibody. (F) Decreased TFII-I SUMOylation in HBE after CBX4 knockdown. HBE was subjected to IP with anti-TFII-I followed by immunoblot analysis with anti-SUMO 1 antibody after CBX4 knockdown. (G) Colocalization of SUMO1 and TFII-I in HDM-treated HBE was determined by immunofluorescence staining of SUMO1 and TFII-I. (H) Co-localization of SUMO1 and TFII-I in HBE transfected with siCBX4 was determined by immunofluorescence staining of SUMO1 and TFII-I. (I) ChIP assay with anti-TFII-I antibody in HBE transfected with the indicated plasmids for 48 hr. Precipitated DNAs were quantified by RT-PCR for promoter regions of MEX-3B gene. (J) Western blots and fold-change of relative luciferase activity against lane 1 in human 293T cells transfected with pcDNA(+)-TFII-I and WT or CBX4 mutants, firefly luciferase reporter, and pRL-tk-renilla plasmids for 24 hr. Data are presented as mean ± SEM. Images show representative results for one of 3 or more experimental replicates. NS, Not significant. (A) Unpaired two tailed Student’s t-test was used. (B, C, D, I and) One-way ANOVA with Bonferroni’s post hoc test was used. \**p* < 0.05 and ** *p* < 0.01.

## Discussion

For the first time, we observed that SUMOylation was enhanced in HDM-induced allergic asthma epithelium and that inhibiting the SUMOylation E2 enzyme reduced airway inflammation, mucus production, and airway hyper-reactivity. Furthermore, we observed that inhibition of SUMOylation significantly decreased the expression of lfTSLP, but not IL-25 and IL-33, in epithelial cells. These results indicate that SUMOylation participates in lfTSLP-mediated allergic airway inflammation.

TSLP is reported to be involved in the regulation of inflammatory processes occurring at the barrier surfaces. For example, a significant upregulation was observed in asthma, atopic dermatitis and *ulcerative colitis*(***Jariwala, et al.,2011***; ***Mitchell and O’Byrne,2017***; ***Park, et al.,2017***). Harada et al provided evidence for the existence of two different isoforms (long form and short form) of TSLP in human bronchial epithelial cells(***Harada, et al.,2009***). It has been reported that the expression of lfTSLP is upregulated while expression of sfTSLP is unaffected in airway epithelium challenged with poly(I:C) and HDM(***Dong, et al.,2016***; ***Harada, et al.,2009***). However, the exact mechanism of this difference in expression remains unclear. Previous studies focused on the difference of their gene promoters (SNP and transcription factors)(***Fornasa, et al.,2015***; ***Harada, et al.,2009***). However, little is known about their post-transcription regulation. In our study, we identified a novel post-transcriptional modification mechanism that specifically regulates lfTSLP, but not sfTSLP, expression. We observed that the SUMOylation E3 ligase, CBX4, can promote lfTSLP expression but has no effect on sfTSLP expression. Unexpectedly, we found that CBX4 does not affect lfTSLP mRNA expression. We subsequently identified that the RNA binding protein MEX-3B, which is regulated by CBX4, can specifically binding to lfTSLP, but not sfTSLP, mRNA and promote its translation.

MEX-3B is a member of MEX-3 family (MEX-3A, MEX-3C, and MEX-3D). This family of proteins binds to specific mRNAs and regulates the expression of their proteins through their two K homology (KH)-type RNA-recognition domains(***Buchet-Poyau, et al.,2007***). MEX-3B has been shown to mediate post-transcriptional stability of IL-33 through its association with the IL-33 mRNA 3 ’ UTR and avoid its degradation in IL-33 induction of ovalbumin allergic asthma model(***Yamazumi, et al.,2016***). Interestingly, we could not detect upregulation of IL-33 in airway epithelium in our HDM-induced allergic asthma and found that MEX-3B regulates lfTSLP RNA translation rather than affecting its stability. Moreover, database prediction suggested that MEX-3B might bind to the lfTSLP mRNA 5’UTR. These contradictory results may be explained by differences in asthma models. These results indicate that MEX-3B exerts multiple post-transcription functions according to the asthma model.

Mechanistically, we identified CBX4 as a transcriptional coactivator of MEX-3B. Despite CBX4 knockdown resulting in the downregulation of MEX-3B mRNA and protein expression, it did not bind to the MEX-3B promoter directly. Although we did not observe an interaction between CBX4 and TFII-I, a transcription factor of MEX-3B, CBX4 enhanced the level of SUMOylation of TFII-I, which promoted its transcriptional activity. TFII-I has been reported to be involved in an array of human diseases, including neurocognitive disorders, systemic lupus erythematosus (SLE), and cancer(***Roy,2017***). To our knowledge, this report is the first to demonstrate a role for TFII-I in TSLP-mediated allergic inflammation. However, several limitations of this study warrant discussion. For example, we failed to obtain samples from patient with asthma for this study. The expression of CBX4 in the airway epithelium of patients with asthma (including those with different inflammation phenotypes) merit additional study. In addition, further investigation of the airway epithelium from a CBX4 KO mice asthma model is warranted.

Collectively, our findings have identified a CBX4/TFII-I/MEX-3B/lfTSLP axis involved in lfTSLP-mediated allergic airway inflammation, suggesting that substrates targeting SUMO E3 ligase activity of CBX4 would be a novel target for the treatment of asthma.

## Methods

### Animals

All animal experimental protocols were approved by Animal Care and Use Committee of Southern Medical University. C57BL/6 mice at 6 weeks of age were used to established the model of asthma. Briefly, mice were sensitized with intraperitoneal 100 μl house dust mite(ALK-Abello A/S, A4963) in 40000U/ml on days 1 and 7, then challenged twice a week by intranasal (i.n.) instillations of 100 μl HDM for a total of seven weeks. Control group received i.n. instillations of PBS alone. For the 2-D08 (MCE, HY-114166) treatment group, 3 mg/kg 2-D08 was administered via oral gavage 2h before every challenge. Every group received equal dosage vehicle (DMSO). Assessments were performed 24 hours after the last i.n. challenge.

### Assessment of airway hyperactivity (AHR), serum lgE and analysis of bronchoalveolar lavage fluid (BALF)

Twenty-four hours after the last challenge, mice airway resistance was performed after challenge with increasing doses of methacholine (0, 3.125, 6.75, 12.5, 25, 50 mg/ml) under anesthesia and mechanical ventilation. The airway resistance at each graded concentration of methacholine was expressed as percentage of baseline value.

After airway resistance measurement, mice were sacrificed with overdose anesthetic. Blood samples were collected by enucleating eyeballs. Blood samples were centrifuged and supernatants were stored at -80 °C. Serum total lgE was quantified by ELISA (RayBiotech).

After blood was taken, bronchoalveolar lavage was performed with 0.5 mL PBS twice and the recovered fluids were pooled. Bronchoalveolar lavage fluid (BALF) was centrifuged at 1500 rpm for 10 mins and the supernatant was used to measure inflammatory cytokines (IL-4, IL-5, IL-13, γ-INF) by LUMINEX multi-factor detection (MERCK). The cell pellets were fixed in 4% formaldehyde for Wright-Giemsa staining and total cells were counted and classified in each slide.

### Histology

The lung tissue was fixed in 4% formaldehyde and embedded in paraffin followed by cutting into sections for hematoxylin and eosin (HE), Periodic Acid-Schiffstain (PAS), immunocytochemistry (IHC) stainning. Sections were deparaffinized with xylene and hydrated with a gradient of ethanol. HE stainning was used to assessed airway inflammatory and PAS was used to assessed mucus production. For IHC, antigen retrieval was carried out through boiling with citrate buffer (pH 6) in a microwave oven for 20 min. Inactivation of endogenous peroxidase was performed by incubating with 30% hydrogen peroxide for 10 min prior to incubating with primary antibody at 4 °C overnight and then with secondary antibody 20 min at room temperature before staining with DAB and counterstaining with hematoxylin. IHC stainning relative average optical density(AOD) in epithelium was analyzed by Image-pro plus 6.0(Figure S4A-H).

### Cell culture, reagents, transfection

Human bronchial epithelial cell line HBE-135-E6E7 (Fuheng biology) was used in this study. HBE were cultured in keratinocyte medium (ScienCell) in a 37°C incubator with 5% CO2 atmosphere. HBE was treated with 2-D08 (5, 10, 20μM) or UNC3866 (MCE, HY-100832) (50, 100, 200 nM) for 24 hours, then 400U/mL HDM (ALK-Abello A/S, A4963) was added for an additional 24 hours. Lipofectamine 3000 (Thermo Fisher Scientific) was used to transfect all siRNAs and plasmids according to manufacturer instructions. The siRNA sequences are listed in supplementary materials (Supplement Table 1).

### Plasmids

pSIN-EF2-PURO-CBX4, pSIN-EF2-PURO-CBX4 (SIM 1/2 mutant), and pSIN-EF2-PURO-CBX4 (chromodomain mutant) plasmids were gifts from Prof. Tie-Bang Kang (Sun Yat-sen University). The plasmids Flag-MEX-3B, Flag-MEX-3B mut KH (G83D, G177D) and Flag-TFII-I were cloned into pcDNA3.1 vector. CBX4 and MEX-3B were knock down by short hairpin. The targeting sequence: CBX4:5’-GCAAGAGCGGCAAGUACUATT-3’; MEX-3B: 5’-CAAUAACAAUAACGGCAAUTT-3’.

### Real-time quantitative RT-PCR

Total RNA was isolated following the protocol by the Trizol kit (TAKARA). SYBR Green (Roche) was used to perform Quantitative RT-PCR by Real-Time PCR instrument (Bio-Rad). The primers were searched on NCBI and listed in supplemental material (Supplement Table 2). The data was calculated using the 2-ΔΔCt method to compare the difference.

### Immunoblot analysis and immunoprecipitation

For Western blots, cells were lysed in RIPA buffer for 15 min, then centrifugated at 14000 rpm at 4 °C for 15 min. 1x SDS loading buffer was added to supernatants and boiled for 10 min. Proteins were separated by SDS-PAGE and transferred to PVDF membranes. The PVDF membranes were blocked with 5% BSA and incubated with primary antibody at 4 °C overnight, then secondary antibody was added at room temperature for 2 hours. The protein bands were detected on an Odyssey imaging system. For immunoprecipitation, cells were lysed in IP lysis buffer (containing protease inhibitor, PMSF, and 20 mM Nethylmaleimide) for 15 min on ice and centrifuged at 14000 rpm at 4 °C for 15 min. The supernatants were precleared using protein A/G beads and the IP primary antibody or negative IgG was added to lysates at 4 °C overnight in rotation before incubating with 50ul protein A/G beads at room temperature for 2 hours. The immuno-complex was collected by centrifugation at 3000 rpm for 2 min and washed three times with 1 mL IP wash buffer. The complex was analyzed by immunoblotting. Antibodies used in this study were as follows: anti-CBX4 (abclonal, A6221), anti-PIAS1 (proteintech, 14242-1-AP), anti-TSLP (abcam, ab188766), anti-IL-25 (abclonal, A8252), anti-IL-33 (abclonal, A8096), anti-MEX-3B (santacruz, sc-515833), anti-FLAG (proteintech, 20543-1-AP), anti-β-actin (proteintech, 60008), anti-sumo1 (santacruz, sc-5308), anti-sumo 2/3 (santacruz, sc-393144), anti-TFII-I (CST, 4562).

### Immunofluorescence

Cells were fixed in 4% PFA for 20 min. Then incubating with primary antibody at 4 °C overnight following incubating with Alexa 488-labeled goat anti-rabbit or 596-labeled goat anti-mouse IgG antibody at 37 °C for 2 hours. Nuclei were counterstained with DAPI for 5 mins. Images were acquired by using a confocal fluorescence microscope (Olympus, Japan).

### Polyribosome Profile Analysis

Sucrose gradient fractionation was carried out as described previously(***Gu, et al.,2009***). Briefly, cells were treated with 100 µg/mL cycloheximide (MCE, HY-12320) at 15 to 30 min prior to harvesting. The cells were lysed with polysome lysis buffer, then centrifuged at 10,000 *x g* for 20 min at 4 °C. The supernatant was added to the top of a 10-50% sucrose gradient. The gradients were centrifuged for 90 min in a SW41Ti swinging bucket rotor at 190,000 *x g* (∼39,000 rpm) at 4 °C and 12-1 mL fractions were collected by upward replacement. The fractions were subjected to RNA isolation and Real-time RT-PCR mentioned above.

### RNA immunoprecipitation assay

RNA immunoprecipitation assay was carried out according to the Magna RIP RNA-Binding Protein Immunoprecipitation Kit manufacturer’s instruction (Millipore). Briefly, cells were lysed in RIP lysis buffer for 15min and centrifuged at 14,000 *x g* for 15 min at 4 °C. The supernatants were collected and immunoprecipitated with antibody to the protein of interest with protein A/G magnetic beads. Magnetic bead bound complexes were immobilized with a magnet and unbound material was washed off. Extract RNAs and detect target gene by RT-PCR.

### Chromatin immunoprecipitation assay

Chromatin immunoprecipitation assay was carried out according to SimpleChIP® Enzymatic Chromatin IP Kit (CST, 9003). Briefly, cells were fixed with formaldehyde to cross-link histone and non-histone proteins to DNA. Then chromatin was digested with micrococcal nuclease into 150-900 bp DNA/protein fragments. Next, antibodies (TFII-I, H3, and lgG) were added and the complex co-precipitated and was captured by Protein G Agarose or Protein G magnetic beads. Finally, the cross-links were reversed, and the DNA was purified and ready for RT-PCR analysis. The CHIP primers are listed in the supplemental material (table 2).

**Table 1.**
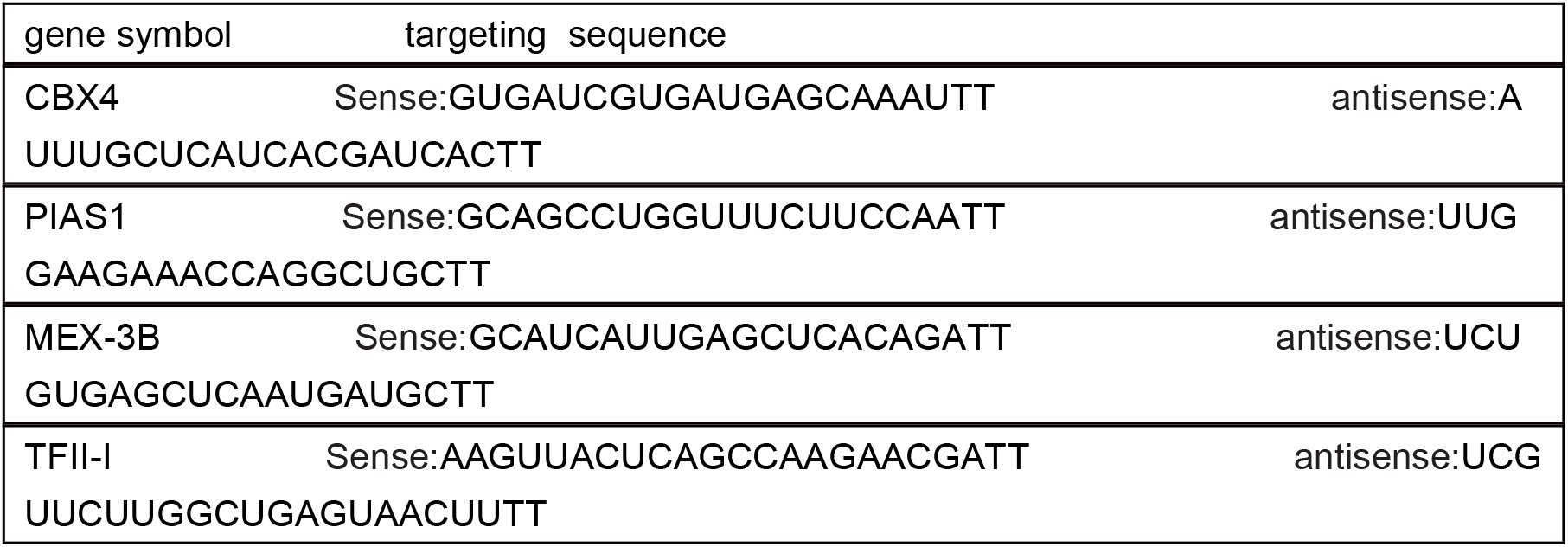
siRNA targeting sequences

**Table 2.**
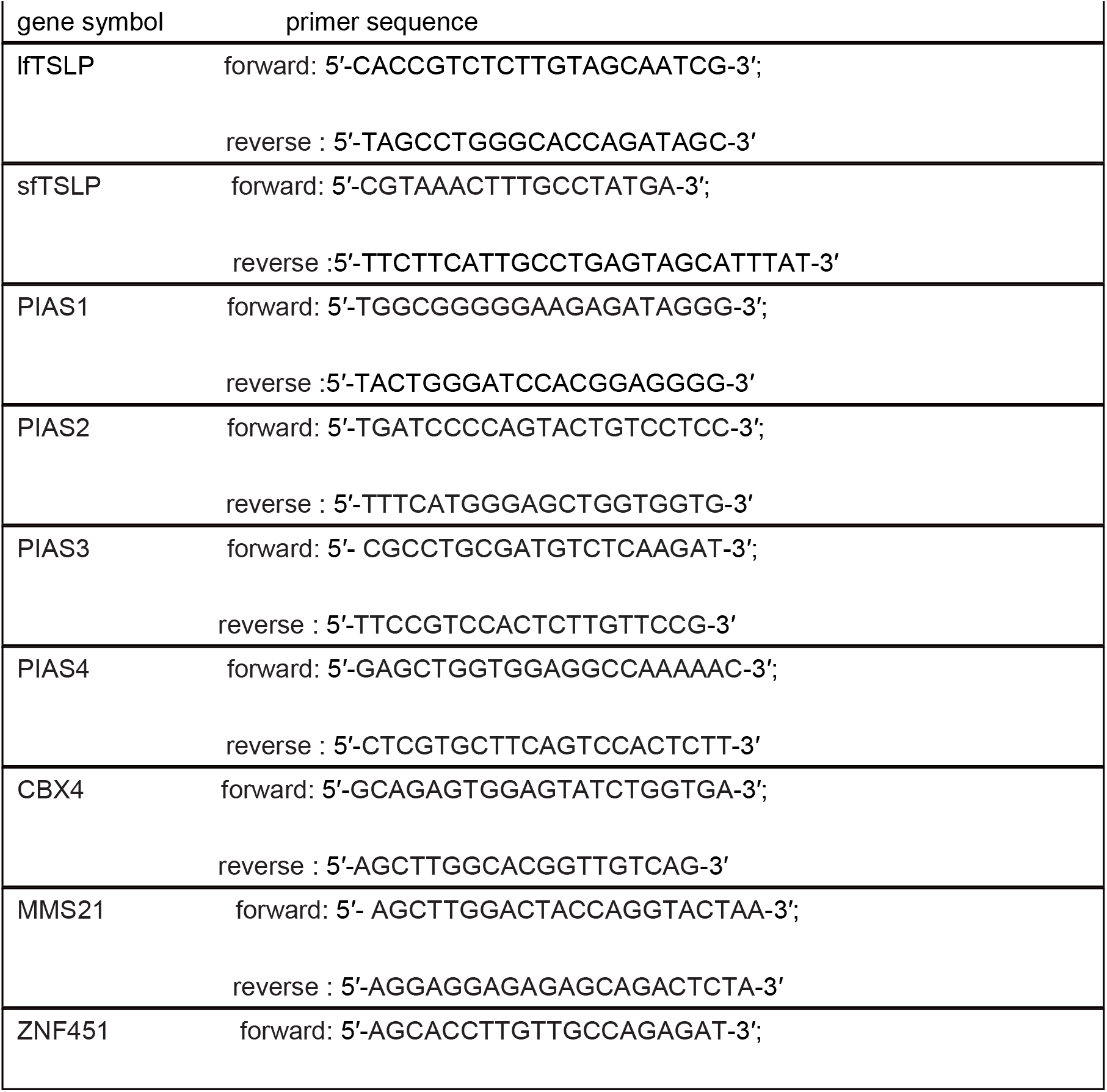
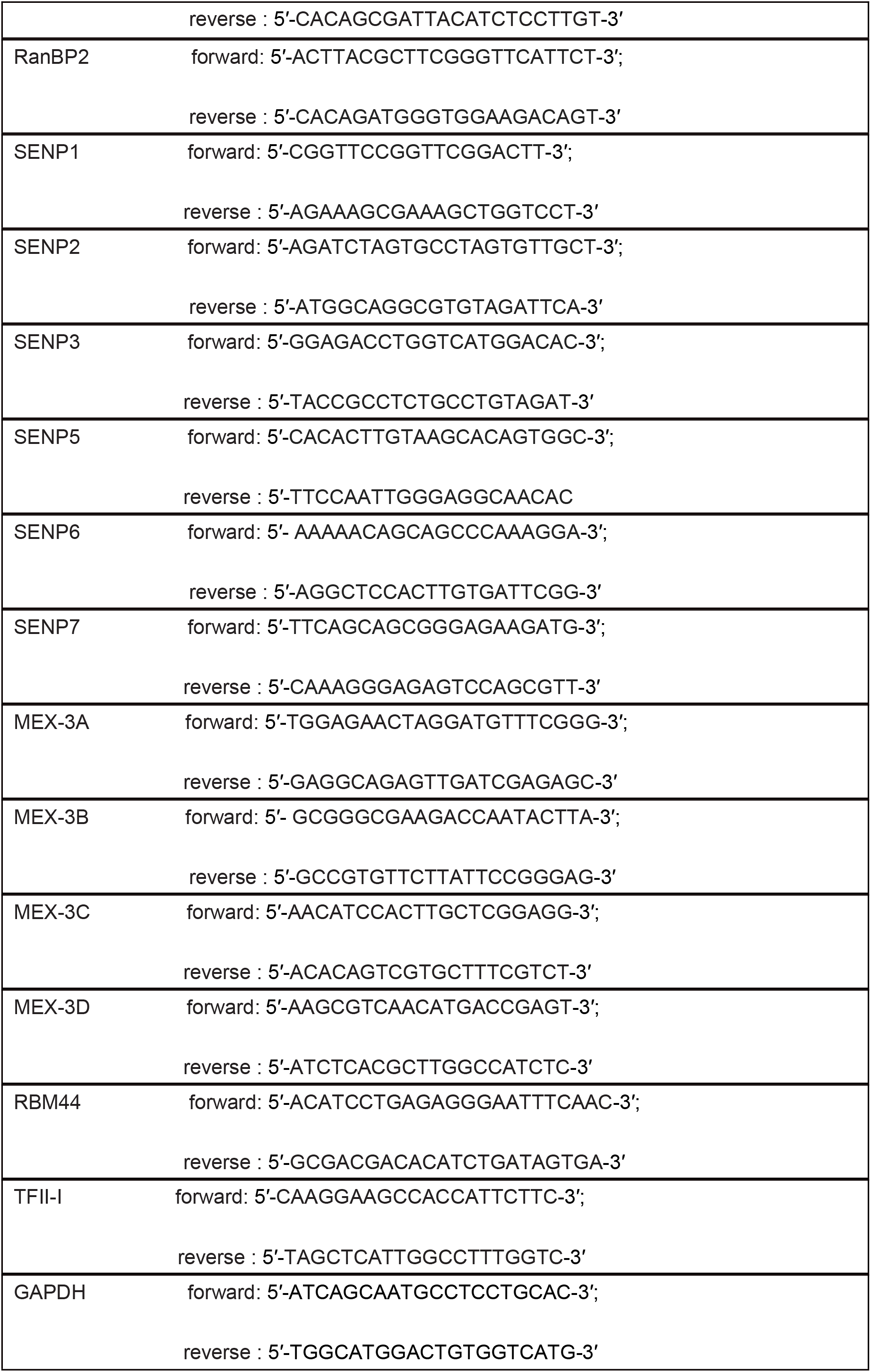
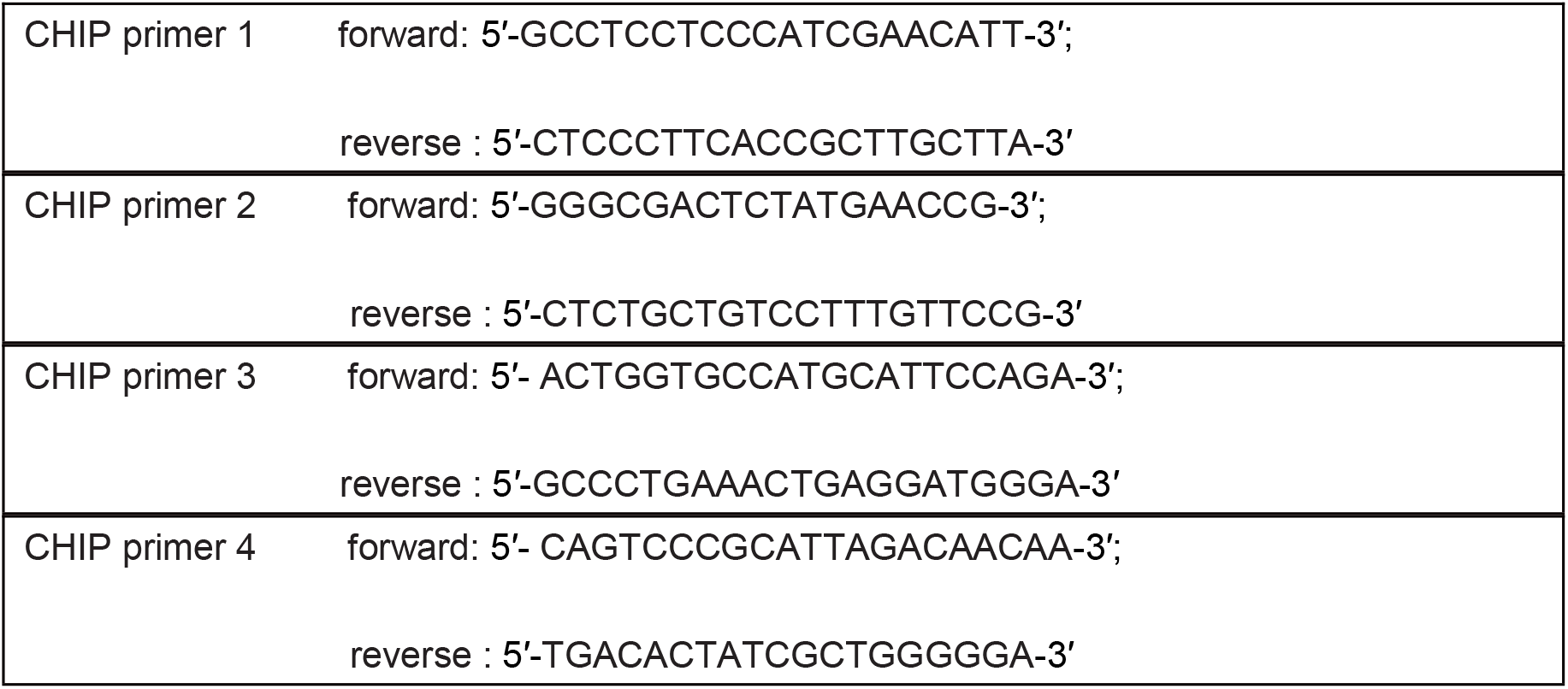
Primers used for qPCR

### Statistical analysis

When comparing two groups, statistical analysis was performed by unpaired two tailed Student’s t-test (normal distribution data). Multiple comparisons were analyzed through one-way ANOVA followed by Bonferroni’s test. AHR data were analyzed using two-way ANOVA. Data were analyzed with GraphPad Prism 6.0 (GraphPad software). The data were presented as mean ±SEM. Differences were considered statistically significant when the *p*-value < 0.05.

## Acknowledgments

We thank Prof. Tie-Bang Kang (Sun Yat-sen University) for pSIN-EF2-PURO-CBX4, pSIN-EF2-PURO-CBX4 (SIM 1/2 mutant), and pSIN-EF2-PURO-CBX4 (chromodomain mutant) plasmids.The authors thank AiMi Academic Services (www.aimieditor.com) for English language editing and review services. This study was supported by the National Natural Science Foundation of China (81700034, 81670026, 8147022, 81770033, 81970032), Natural Science Foundation of Guangdong Province (2017A030310106, S2013040013505, 2014A030310325, 2015A030313236, 2014A030310325, 2017A030313849), the Precision Medicine Research of The National Key Research and Development Plan of China (2016YFC0905803), and China Postdoctoral Science Foundation (2017M620377, 2018T110885). The authors declare that they have no conflicts of interest.

## Abbreviations

CBX4: chromobox 4
HDM: house dust mite
lfTSLP: long-form thymic stromal lymphopoietin
SUMO1: Small Ubiquitin Like Modifier 1.

## Graphical Abstract

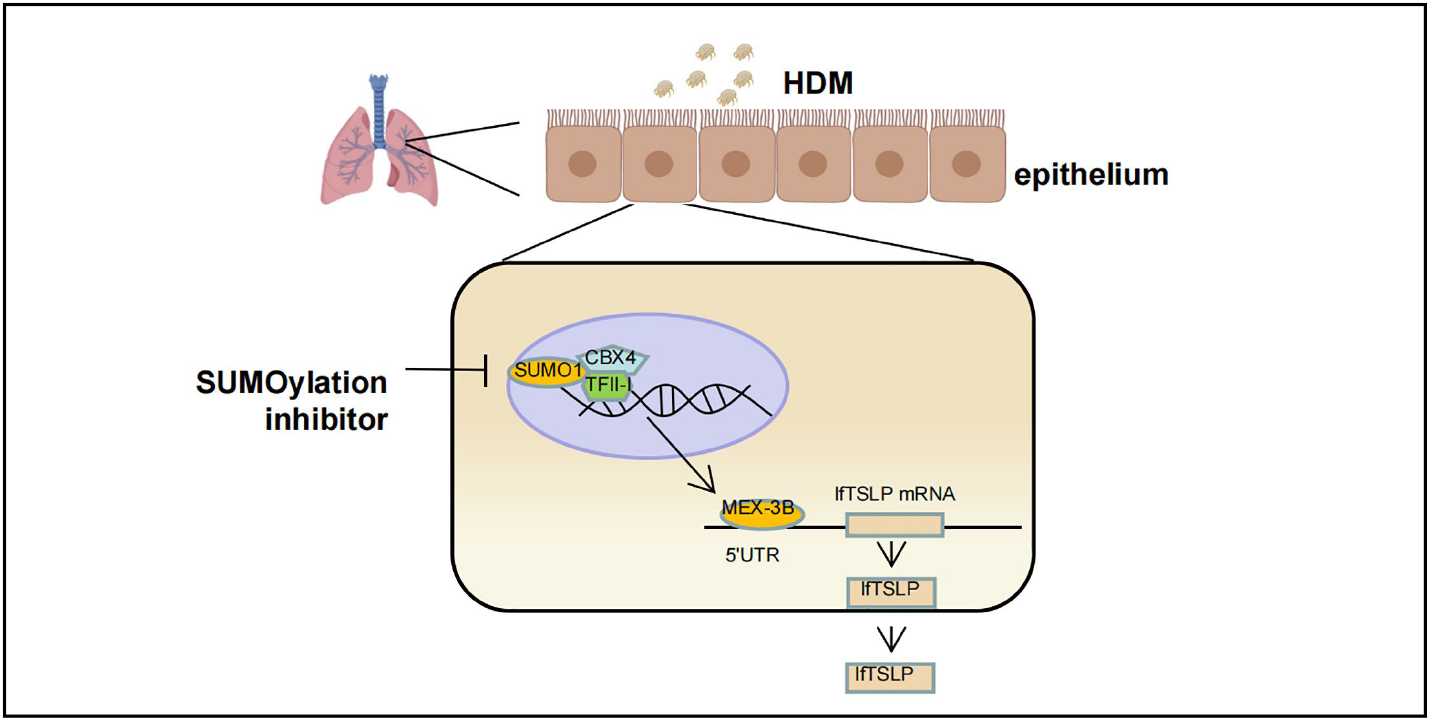

## Supplement

**Figure S1.**
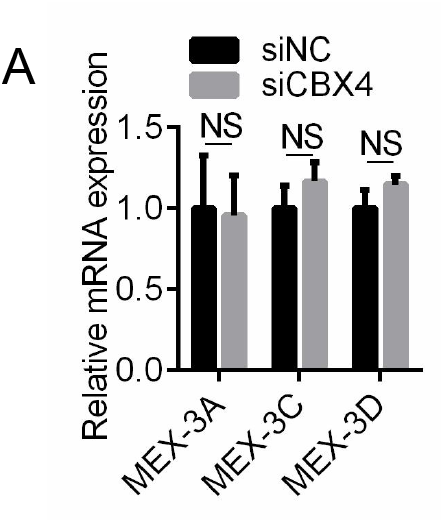
MEX-3A, MEX-3C and MEX-3D mRNA levels were detected by qRT-PCR after knocking down CBX4 in HBE. Unpaired two tailed Student’s t-test was used. NS, Not significant.

**Figure S2.**
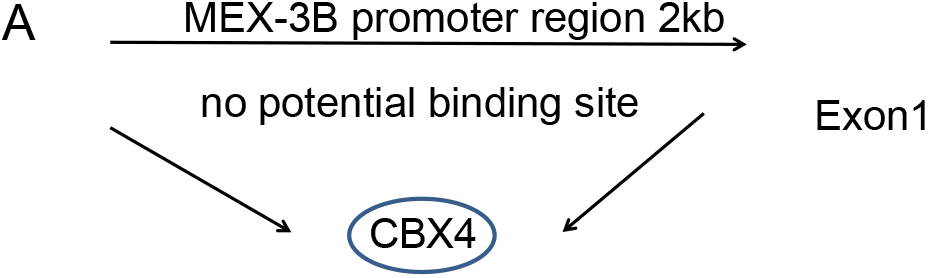
The potential interaction of CBX4 and MEX-3B promoter was scanned in TFtarget database.

**Figure S3.**
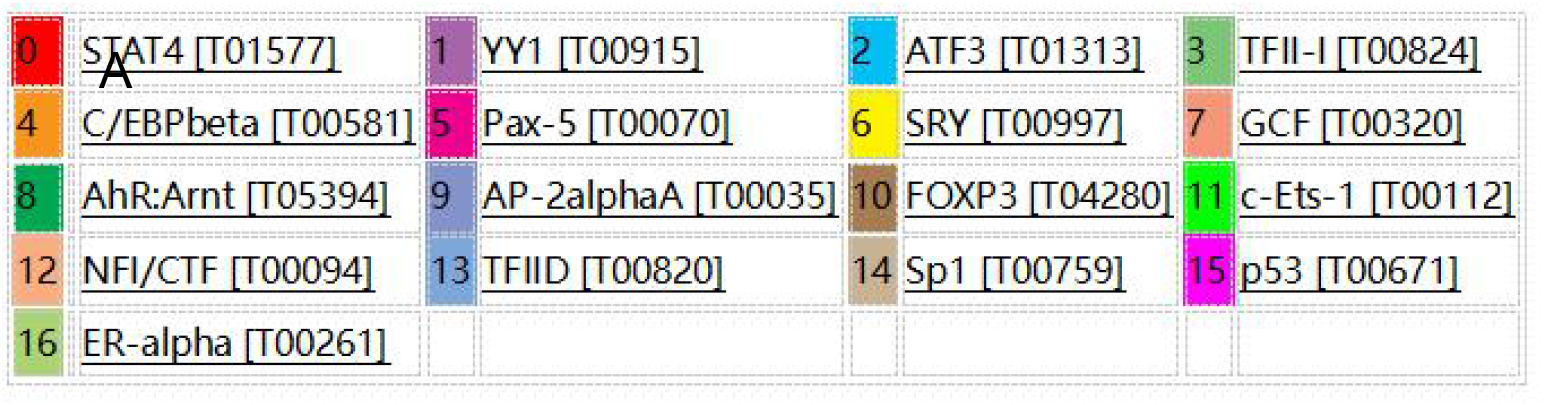
The potential transcription factors of MEX-3B was scanned in PROMO database.

**Figure S4.**
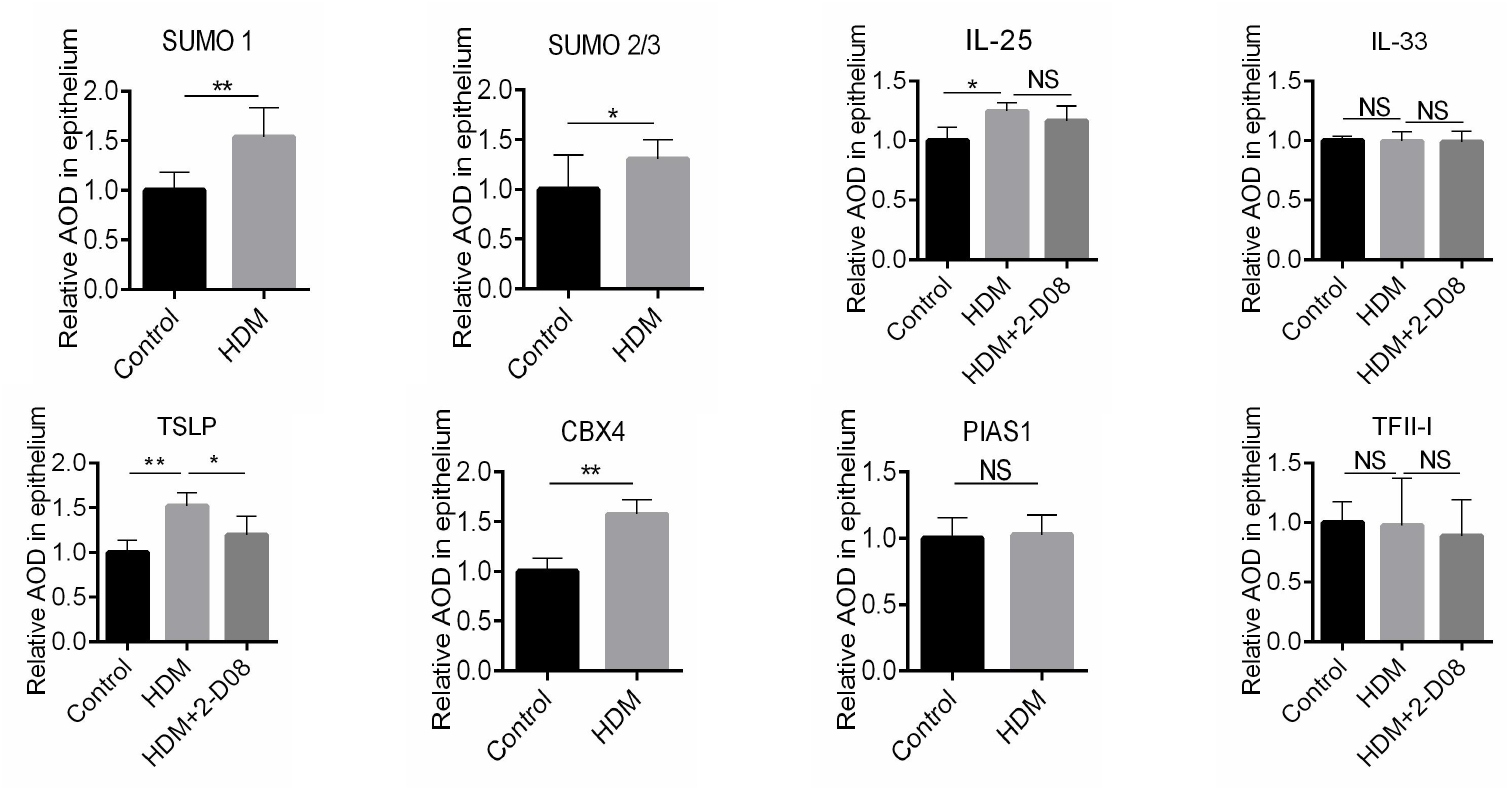
Quantitative immunohistochemical analysis was performed for SUMO 1(A), SUMO 2/3(B), IL-25(C), IL-33(D), TSLP(E), CBX4(F), PIAS1(G), TFII-I(H) in airway epithelium. Data are presented as mean ± SEM. Images show representative results for one of 3 or more experimental replicates. NS, Not significant. (C, D, E and H) One-way ANOVA with Bonferroni’s post hoc test was used. (A, B, F and G) Unpaired two tailed Student’s t-test was used. * p < 0.05, ** p < 0.01, and *** p < 0.001.

